# Fresh Snowfall Microbiology and Chemistry are Driven by Geography in Storm-Tracked Events

**DOI:** 10.1101/300772

**Authors:** Honeyman A. S., Day M.L., Spear J.R.

## Abstract

Snowfall is a global phenomenon highly integrated with hydrology and ecology. Forays into studying bioaerosols and their dependence on aeolian movement are largely constrained to either precipitation-independent analyses or *in*-*silico* models. Though snowpack and glacial microbiological studies have been conducted, little is known about the biological component of meteoric snow. Through culture-independent phylogenetic and geochemical analyses, we show that the geographical location at which snow precipitates determines snowfall’s geochemical and microbiological composition. Storm-tracking, furthermore, can be used as a valuable environmental indicator to trace down what factors are influencing bioaerosols. We estimate annual deposits of up to ~10 kg of bacterial / archaeal biomass per hectare along our study area of the eastern Front Range in Colorado. The dominant kinds of microbiota captured in an analysis of seven snow events at two different locations, one urban, one rural, across the winter of 2016/2017 included phyla *Proteobacteria*, *Bacteroidetes*, *Firmicutes* and *Acidobacteria*, though a multitude of different kinds of organisms were found in both. Taxonomically, *Bacteroidetes* were more abundant in Golden (urban plain) snow while *Proteobacteria* were more common in Sunshine (rural mountain) samples. Chemically, Golden snowfall was positively correlated with some metals and anions. The work also hints at better informing the ‘everything is everywhere’ hypotheses of the microbial world and that atmospheric transport of microbiota is not only common, but is capable of disseminating vast amounts of microbiota of different physiologies and genetics that then affect ecosystems globally. Snowfall, we conclude, is a significant repository of microbiological material with strong implications for both ecosystem genetic flux and general bio-aerosol theory.

**Importance:** Snowfall is commonplace to the temperate and polar regions of the world. As an interface between the atmosphere, hydrosphere and earth, snow is responsible for high annual deposits of moisture globally, and, can serve as a ‘water bank’ in the form of both permanent snow fields and glaciers. Essential to general ecosystem function, snow can also be considered a transporter of aerosolized material. Given the magnitude of microbiota deposited by snowfall, which we report, it is likely that biological material within snowfall, with its geochemical underpinning— and the associated genetic banks—have significant downstream ecological effects.

Understanding what is contained in snowfall becomes especially urgent in a warming climate where high-impact meteorological and ecological changes are imminent and likely. With climate-induced changes to snowfall patterns, surface ecosystems are likely to be impacted by ensuing changes in microbiota deposition. Thus, the ecosystem function of soils, rock and surface waters are also likely to be impacted; these changes, in turn, greatly influence agriculture, weathering and infrastructure.

## Introduction

Throughout temperate and polar regions of the world, snowfall is ubiquitous. Specifically, along the Front Range of eastern Colorado, the mean annual snowfall from 2010-2016 is reported as 97.7 inches in Boulder, Colorado (1). Despite snow’s prevalence, little is known about the biological composition of this massive annual hydrological deposit; meteoric snow is an essential part of yearly moisture input in Boulder, Colorado given that it represents 43% of the total precipitation (by liquid water volume) from 2010-2016 (1). Both snow and rain require an initiation surface for atmospheric water to condense into a droplet of water or ice particle, respectively. Typically, these are thought to be airborne particles of particulate matter, both organic and inorganic of various colloidal sizes (e.g., 0.5 to 8 μm) (2, 3). Bioaerosols, defined as particles of a biological nature released from both terrestrial and marine ecosystems into the receiving atmosphere, are considered to be capable of acting as initiation surfaces (4). Recently, bioaerosols have also been suggested as an understudied component of both atmospheric processes and biogeographical fate (5).

Bioaerosols are most famous for their potential detrimental effects upon human health from contaminated indoor or other built-environments (e.g., mold and fungal spores) (6). However, even in a benign, typical, outdoor / atmospheric environment, bioaerosols are common and the literature reports source environments (7), seasonal effects (8, 9), and diurnal shifts (10) as factors in atmospheric bioaerosol composition. Furthermore, bioaerosols are known to be components of the estimated 3 Pg annual dust traffic around the globe (11–13). Work has approached the theory that bioaersols are important to climate as a whole (14) by examining *in*-*silico* models of microbiological dispersion in the atmosphere (15). With respect to precipitation, the effects of rainfall additions to soil communities has already been shown to be of microbiological community significance at the soil / atmosphere interface (16). Additionally, these works suggest that snowfall, in particular, is even more important in the deposition of some strains of ice-nucleating bacteria *Pseudomonas syringae* (also a plant pathogen) (17)—from which the man-made snow-making additive Snomax^™^ derives (18). *In*-*vivo* studies of both atmospheric and hail microorganisms have shown them to be metabolically active (19–23), suggesting that the biomass in the environment from which snowfall derives is not of inconsequence. In addition, microbiota can remain metabolically active throughout the rain / ice / snow nucleation process. What microbiota within rain / ice / snow precipitation look like when they contact soil and surface receiving waters, however, remains unknown. Beyond the community structure of precipitation microbiota, it is especially unknown how these structures changes as a result of geography and atmospheric events.

Bioaerosol literature, in general, is best represented by reports of *in*-*silico* bioaerosol movement, aerosol microbial composition and studies of canonical surface constituents such as glaciers and snowpack (24–28). There exists little specific information, however, on the microbiology of snowfall precipitation. The biological composition of snowfall—a massively spatial meteorological event—should be well understood vis-à-vis magnitude and community structure, especially given that precipitation is key to linking atmospheric and surface theory. An aid to such study is the preponderance of weather information available that tracks storm events in multiple-spectral analyses; these data further inform storm trajectories as well as hydrologic source-water locations.

To this end, we sampled fresh snowfall throughout the entirety of the 2016-2017 snow season along the Front Range of eastern Colorado. Our investigations include disparate sampling sites (one rural mountain, and one urban plain) to help in the characterization of snowfall with respect to both sampling location—within the same storm—and direction of origin of the storm. We hypothesized that the biological content of fresh snowfall would be uniquely distinguishable with respect to sampling location and that storm tracking could provide novel insight into other factors that may affect biogeography. Here, we apply ecological investigative approaches together with remote sensing based atmospheric storm analyses to assess the magnitude of biological deposition, community structure, and abiotic solute associations in fresh snowfall.

## Methods

### Sample Collection

Over the course of the 2016-2017 snow season, fresh snowfall samples were collected from two locations along the Front Range of eastern Colorado. One sampling location was a rural setting (Sunshine, CO; latitude 40.060623, longitude -105.365711, elevation 2,164 m) and the other more urban / suburban (Golden, CO; latitude 39.751964, longitude -105.224620, elevation 1,730 m). Throughout the snowfall season, seven snowstorms were sampled. Three of the storms yielded snow at both locations, allowing paired sampling of the storm on the same day. Snow collections were conducted either the evening of a day storm, or the morning after a night storm. Sealed, unopened, Ziploc^™^ bags were used to remove the top ~5-8 cm of accumulated snow at each sampling event. Typically, four one-gallon (3.8 L) bags of snow were collected per sampling event, with the bags contacting neither hands, nor ground beneath the fresh snowfall. When sampling a storm, either there was no prior ground accumulation, or snow was only sampled modestly from the top layer, leaving many inches between the sampling scoop and snowfall from a previous storm. Snow in sample bags was kept frozen with wet ice during transport, and at -20°C during storage. Immediately prior to filtering, snow samples were thawed within their sealed bags via heated water over 10 minutes; under this thawing protocol melted snow water could not exceed 0°C until all snow was melted, at which point the bags were removed from the heated water. Using a sterile 0.45-μm 500 mL Autofil PES Bottle Top vacuum filter cup with lid (Foxx Life Sciences, Salem, NH), thawed snow samples were filtered. Filtrate was collected into autoclaved 1 L glass bottles. Filter papers were sectioned into quadrants, with three used as technical replicates for DNA extraction, and one used for imaging. All quadrants were frozen at -20°C until further processing. As a control for the filtering process, one additional one-gallon bag of snow was collected on 04/04/2017 from Sunshine, autoclaved twice at 121 °C and 15 PSI for 30 minutes in a glass bottle, and then filtered and processed the same as other samples.

### Microscopy

Filter paper quadrants from each sample were imaged on both a Hitachi TM-1000 (Hitachi High Technologies in America, Schaumburg, Illinois) environmental scanning electron microscope (ESEM) equipped with a Bruker (Billerica, Massachusetts) electron dispersive spectroscopy detector as well as an Olympus CX41 phase contrast microscope (Olympus Corporation, Center Valley, Pennsylvania) equipped with an Infinity 2 camera and Infinity Analyze software (Lumenera Corporation, Ottawa, Ontario).

### Storm Tracking

Snow storms associated with sample collection were tracked with the NOAA ARL (Air Resource Laboratory) HYSPLIT Trajectory Model (29). A trajectory ensemble was generated for each storm by creating a set of tracking vectors centered about the latitude and longitude of each sampling site. Models are backward trajectories and encapsulate storm tracking up to 24 hours prior to the day of the snow storm event. Backward trajectories began at 500 m AGL (above ground level) from sampling sites.

### Geochemistry

Filtrate from melted snow was subsampled to two 15 mL aliquots for Ion Chromatography (IC; major anions) and Inductively Coupled Plasma Optical Emission Spectrometry (ICP-OES; major cations) analysis. ICP-OES samples were acidified with three drops of trace metal grade 10% Nitric Acid. IC analysis was performed on a Dionex ICS-90 ion chromatography system running an AS14A (4 × 250 mm) column, while ICP-OES was performed on a Perkin-Elmer Optima 5300 DV Inductively Coupled Plasma Optical Emission Spectrometer.

### DNA Extraction

The ZymoBIOMICS DNA Mini Kit (Zymo Research, Irvine, CA) was used according to manufacturer instructions for DNA extractions. Input into the extraction process was one quadrant of a 0.45-μm filter (quadrants described in ‘Sample Collection’). For every set of extractions conducted, an extraction blank (a DNA extraction conducted with no sample input) was also processed. A single sample replicate from Golden taken on 11/17/16 was removed from all downstream analyses due to microcentrifuge tube failure.

### Quantitative PCR (qPCR) and DNA Sequencing

To quantify the abundance of Bacteria and Archaea in the snow samples,_LightCycler 480 SYBR Green I Master qPCR mix (Roche, Indianapolis, IN) was used in conjunction with 16S rRNA V4-5 region primers (30, 31). Genomic DNA from *Escherichia coli* strain JM109 was used as a standard from a Femto Bacterial Quantification Kit (Zymo Research, Irvine, CA) to calibrate amplification curves; 7 standards, in addition to a no-template (negative) control (NTC), ranged from 2×10^-5^ to 20 ng of DNA per reaction well. qPCR reaction volumes were 20 μL of 1x Master Mix, 0.2 μM 334F (CCA GAC TCC TAC GGG AGG CAG C), 0.2 μM 519R (GWA TTA CCG CGG CKG CTG), and 2 μL volume of template DNA. Amplification was conducted in technical triplicate with a LightCycler 480 II (Roche, Indianapolis, IN). LightCycler amplification parameters were as follows: initial denaturation at 95°C for 5 min (4.4°C/s), 40 cycles of 95°C for 30 s (4.4°C/s) then 53°C for 30 s (2.2°C/s) and 72°C for 30 s (4.4°C/s), final extension of 72°C for 5 min (4.4°C/s).

DNA sequencing was performed on an Illumina MiSeq (Illumina Inc., San Diego, CA) at the Duke Center for Genomic and Computational Biology using V2 PE250 chemistry. Initial sample 16S/18S rRNA PCR amplification was performed in duplicate 25 μL reactions as follows: 1x QuantaBio 5PRIME HotMasterMix, 0.2 μM 515-Y Forward Primer (5’-<u>GTA AAA CGA CGG CCA GT</u> CCG TGY CAG CMG CCG CGG TAA-3’) where the M13 forward primer (underlined) is ligated to the small sub-unit RNA (ssuRNA) specific sequence via a ‘CC’ spacer, 0.2 μM 926R (5’-CCG YCA ATT YMT TTR AGT TT-3’) Reverse Primer and template DNA. Reverse primers were as reported by Parada et al. (2015) (32) while forward primers were adapted from Parada et al. with the addition of the M13 sequence, as done by Stamps et al. (2016) (33). Primers from Parada et al. target all three domains of life (32). PCR was performed on a Techne TC-5000 with the following parameters: initial denaturation at 98°C for 30 s, followed by 30 cycles of 98°C for 10 s then 52°C for 20 s and 72°C for 10 s, with a final extension of 72°C for 5 min and a 10°C hold. A second, 6 cycle, PCR amplification step with the parameters used in the first amplification step was used to ligate barcodes adapted from Caparaso et. al (2012) (34) to the M13 region of the forward primer (33). PCR product cleanup was performed with 0.8x concentration of KAPA Pure Beads (Kapa Biosystems, Indianapolis, IN) according to the manufacturer’s specifications.

### Sequence Analysis

DADA2 (35) was used to trim sequences, infer sample composition, merge Illumina paired-end reads, construct a sequence table, and remove chimeric sequences. Initial trimming of the sequences involved excising forward and reverse primer sequences. Technical replicates from each sample were pooled into one object and analyzed together; these pooled samples will be referred to simply as ‘samples’ from here on. Samples were trimmed to contain only bacterial and archaeal sequences, with chloroplast and mitochondrial sequences similarly excised. All samples were rarefied to a depth of 3,391 sequences; 3,391 was chosen because it allowed inclusion of all but one sample (12/07/16 Sunshine) while also retaining most of the sample diversity (Figure S1). Downstream sequence analyses were run with multiple random subsampling sets and results were not found to change appreciably at our selected rarefaction level of 3,391 bacterial / archaeal sequences. For reproducibility, a set random start seed was used in the rarefaction process. Taxonomy assignments were called with a SILVA training file through the DADA2 taxonomy assignment script. Phylogenetic trees were constructed with sequence alignment via SINA (Silva Incremental Aligner; v1.2.11; 90% complementarity) (36) in the QIIME package (37). OTU tables, taxonomy tables, phylogenetic trees, and sample metadata were compiled in the data processing phyloseq object presented by the R package ‘phyloseq’ (38).

Assessment of eukaryotic sequences was done via the same method as above, with the exception of merging forward and reverse Illumina reads. In a separate pipeline, all forward and reverse reads were merged by concatenating the reads directly with a 10-Ns as the joiner. As in the bacterial / archaeal analysis, sequences from technical replicates were pooled to one sample. Bacterial and archaeal taxonomic assignments were removed from this dataset, leaving only 18S rRNA eukaryotic sequences. The eukaryotic sequence dataset was rarefied to 529 sequences; few eukaryotic sequences were returned in general, and rarefaction at 529 sequences therefore did not dramatically reduce the number of sequences analyzed. Like the bacterial and archaeal analysis, a set random start seed was used during rarefaction for reproducibility. Three samples were removed from the eukaryotic analysis because they were below the sequence threshold (two from Golden and one from Sunshine).

## Results

### Microscopy

Imaged filter quadrants from all samples revealed patterns that distinguish samples taken from Golden from those taken in Sunshine (Figure 1). There is substantially more material captured by filtering snow from Golden than snow from Sunshine in the same storm. Any storm localized to Golden generally yielded more material than storms localized to Sunshine. A large number of morphologically diverse particles were collected from both locations; i.e. sizes of captured matter range from just above the 0.45 μm threshold to 30 μm and greater. Cells are not visible within microscopy images, which was expected given the concentrations of cells that were filtered and their resulting spatial distribution on filter papers.

**Figure 1:**
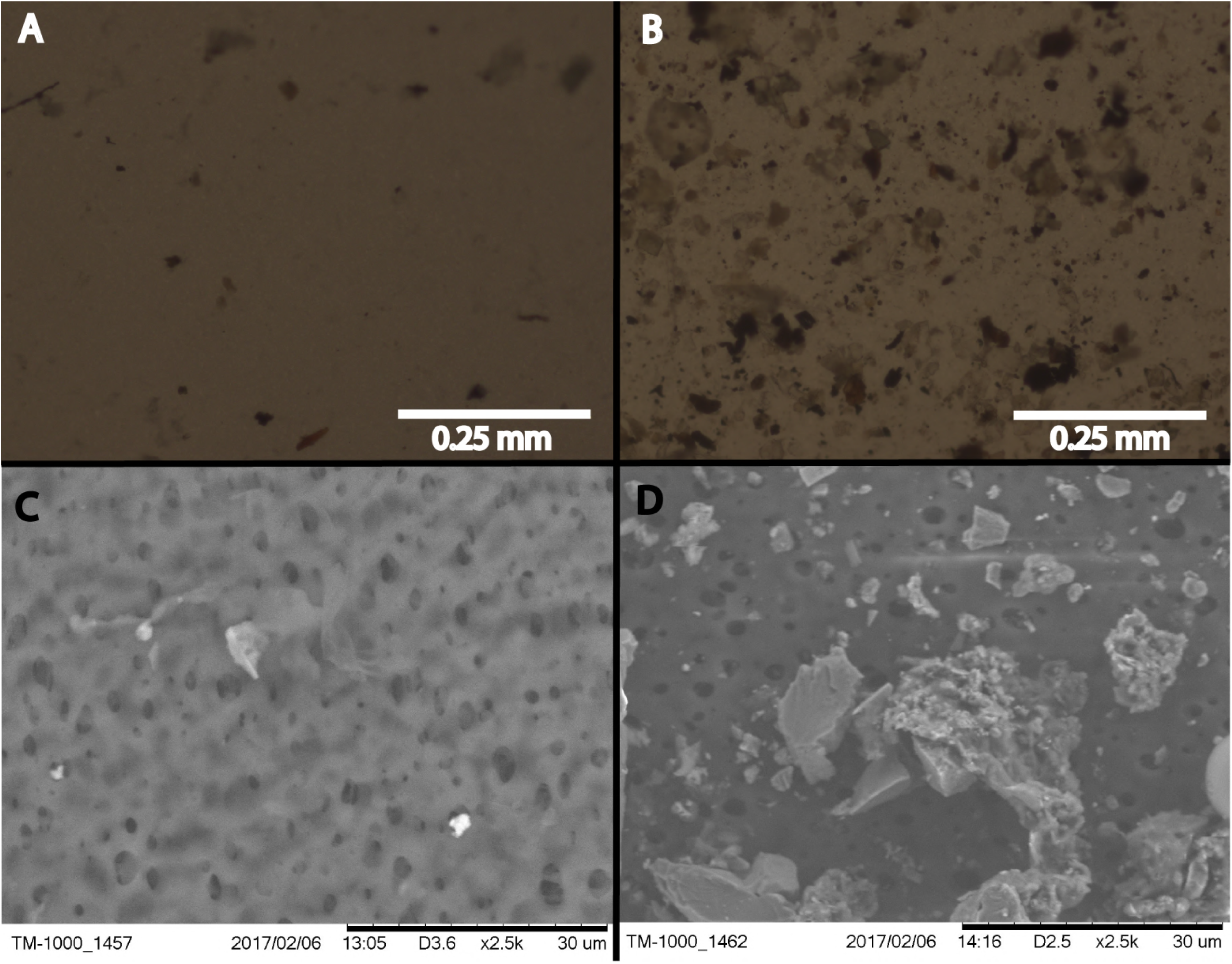
Bright-field (**A and B**) and ESEM (**C and D**) imaging of one quadrant of the filter paper that was used for the snowstorm on 01/04/17. Panels **B** and **D** are the filter paper from filtered Golden snow while panels **A** and **C** show filtered material from Sunshine. All filter papers are rated to capture 0.45 μm, and greater, particles.

### Storm geochemistry

IC and ICP-OES analyte results were compiled to generate a principal component analysis (PCA) (base R stats package) with individual samples as the object of ordination (Figure 2). To reduce noise in the model, only analytes seen to have variation amongst samples were used in the ordination and these were: Cl, NO_2_, NO_3_, SO_4_, Ca (315.887.R), K (766.490.R), Mg (279.553.R & 285.213.R), Mn (257.610.A), Na (589.592.R), S (180.669.A), Si (251.611.A), and Zn (213.857.A). Adjacent to the wavelengths in parentheses, ‘A’ and ‘R’ indicate either Axial or Radial viewing during measurement. Data were centered and scaled to unit variance. Normal ellipses (Figure 2) show nominal clustering by sampling location. Furthermore, the arrows in Figure 2 indicate a positive correlation of variables in the direction that they point. Except for Zn, Mn, and Si, all analyte variables correlate positively with snow collected in Golden. Principal Component 1 suggests a high chemical variance between samples collected in Golden relative to those collected in Sunshine. To assess the statistical differentiation of chemical data with respect to sampling location (independent of the PCA), a Euclidean distance matrix was constructed using the same samples and chemical data used in the chemical PCA analysis. An Adonis test was run on the Euclidean distance matrix to inspect statistical differences between Sunshine and Golden snowmelt chemistry; the two groups were found to be significantly different (R^2^ = 0.28, p = 0.002).

**Figure 2:**
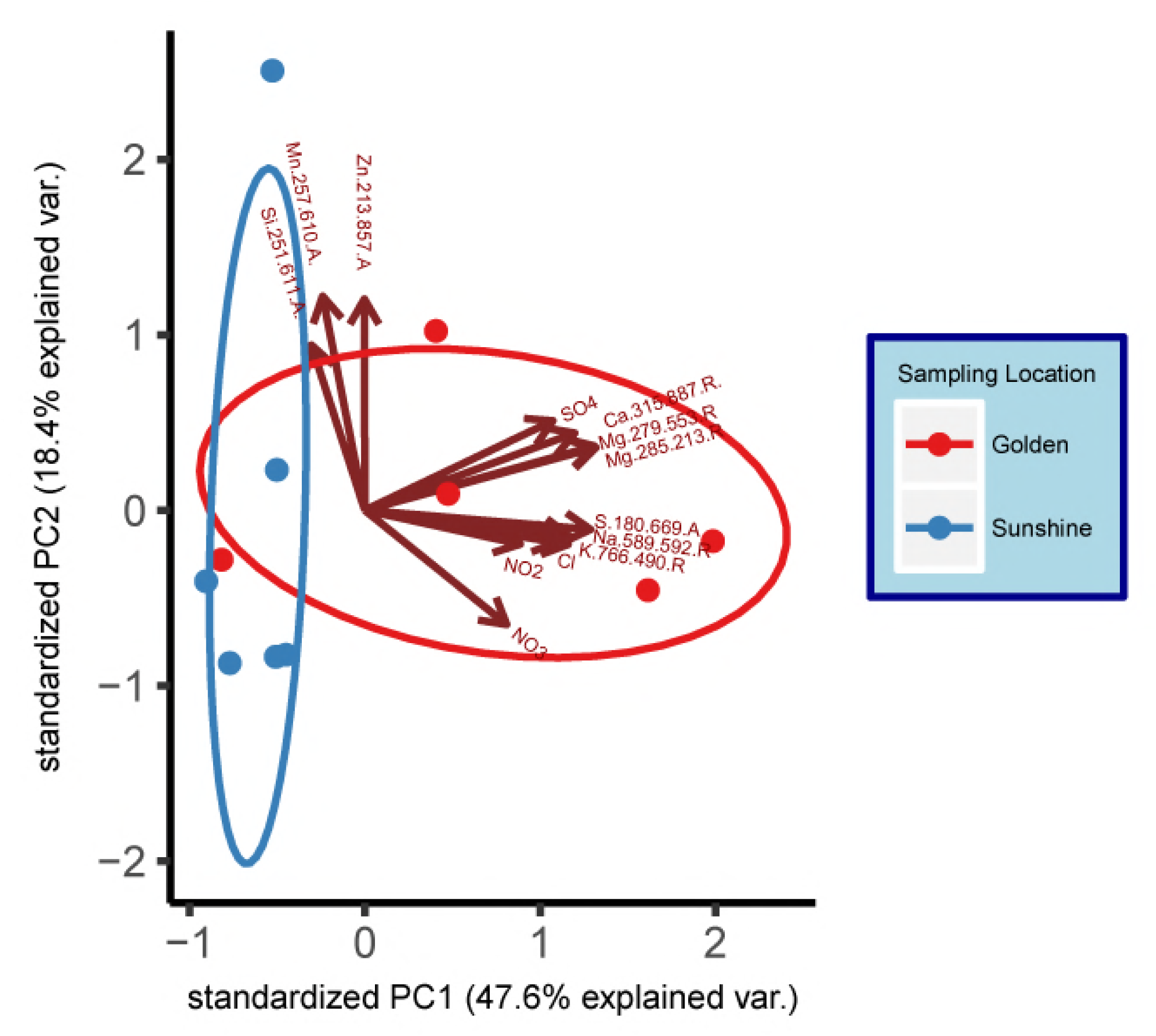
Principal component analysis (PCA) of geochemical data (IC/ICP-OES) grouped by sampling location. Data have been centered and scaled to unit variance. Grouping ellipses are normal data ellipses constructed about data points from each sampling location.

### Genomic Yield and Sample Diversity

16S rRNA gene amplification via qPCR was used to estimate the amount of bacterial / archaeal genomic DNA extracted from each sample. DNA extracted per sample was normalized to the volume of filtrate that was yielded by the filtering process. Reported values determined by qPCR were in units of estimated nanograms of genomic DNA (gDNA) extracted per milliliter of snow filtered (Figure 3). At least one filter replicate from all samples amplified a minimum of 2 qPCR cycles before negative controls; all filter replicates were used for each sample in downstream analyses. We are unable to include the 12/7/2016 Sunshine sample on Figure 3 because we do not have data on the volume of melted water resulting from filtering four one-gallon (3.8 L) bags of snow. When possible, snow collected from the same storm in disparate locations are co-plotted and p-values from a student’s t-test reported (Figure 3). The p-value was significant in one of the three comparisons conducted. However, snow collected from Golden generally yielded higher amounts of genomic DNA than snow collected from Sunshine in the same storm. Across all samples in our study, the average estimated mass of bacterial / archaeal genomic DNA (gDNA) extracted per milliliter of snow-water filtered was 0.237 ng mL^-1^, which (using *E. coli* —~ 5 fg gDNA cell^-1^—as a model), corresponds to a likely cell count of ~4.7 × 10^4^ cells mL^-1^ of snow melt.

**Figure 3:**
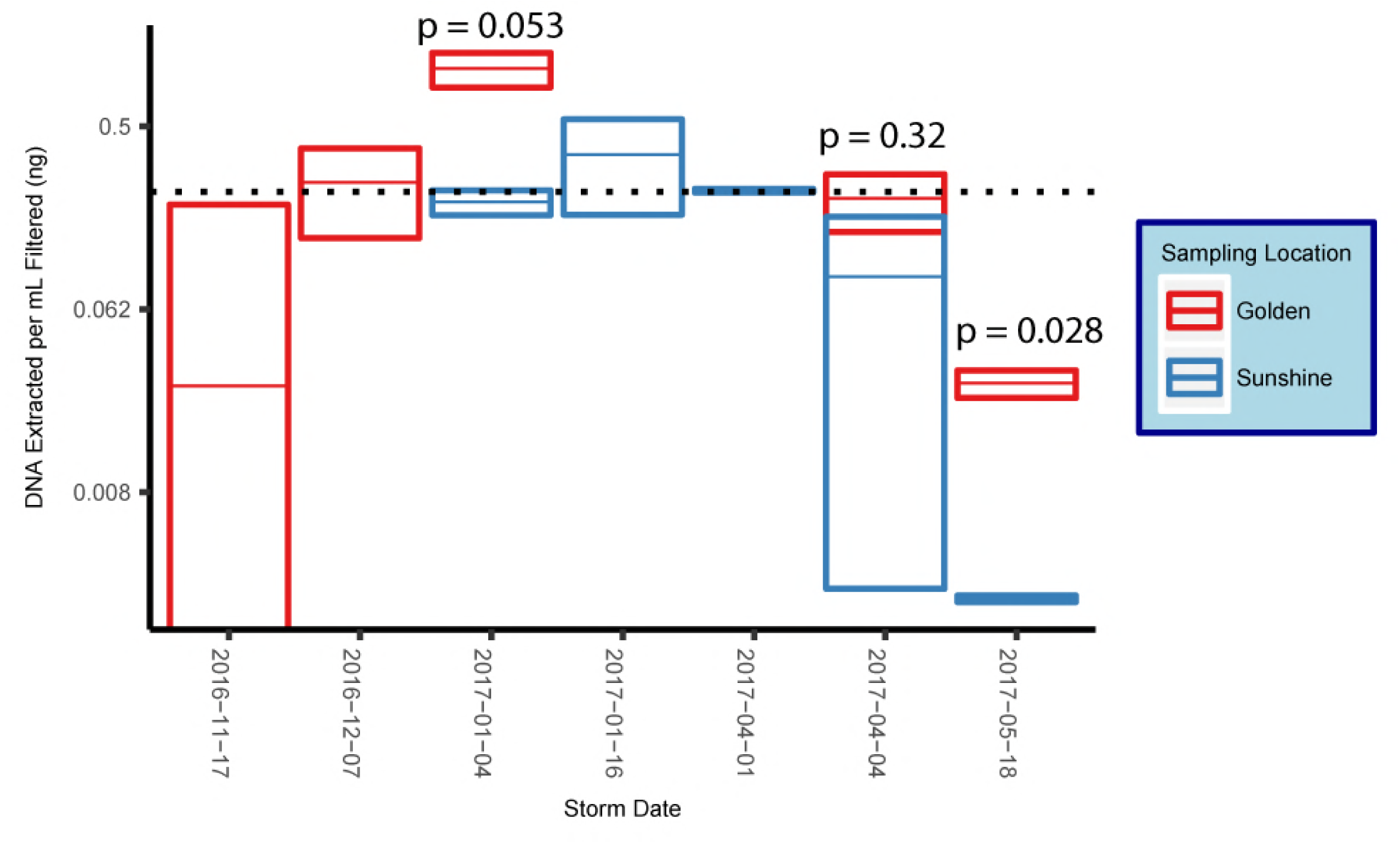
Box plots of mean mass of gDNA extracted from replicate filter quadrants per mL of snow filtered. Error bars are standard errors, indicating technical variance. The dotted line represents the mean value across all samples and all snowstorms for the 2016-2017 snow season along the eastern Front Range in Colorado. Reported p-values are from a student’s t-test.

Post-rarefaction—to a number of bacterial / archaeal sequences where most diversity could be retained and only one sample was cut out (12/07/2016 from Sunshine)—bacterial / archaeal alpha diversities were estimated with the number of observed OTUs. The mean number of observed OTUs for Golden samples was 188.0 while the mean number seen in Sunshine was 98.4 (p = 0.032, Welch two sample t-test). While the alpha diversity of snow sampled in Golden is higher than Sunshine Canyon, the alpha diversities between different snow storm origins have insignificant p-values.

### Storm Tracking

NOAA HYSPLIT models were used to assess which direction the majority of storm travel vectors came from for each sampling event. Most storms were of a SW origin while two storms came from the NW and one storm was of a SE origin (a Colorado ‘upslope’ storm). We verified from the HYSPLIT models that same-day samples from both geographic locations were in fact from the same snow storm and could be assigned the same direction of origin (example: Figures S2 and S3).

### Characterization of Snow Communities

A community ordination chart organized by both sampling location and origin of storm is presented. Principal Coordinate Analysis (PCoA) suggests strong clustering by sampling location, as constructed by a weighted Unifrac distance matrix (Figure 4) (39). An Adonis test was run on the dissimilarity matrix used for Figure 4, and sampling locations were found to be distinct in grouping (R^2^ = 0.20, p = 0.033). Weighted Unifrac distance matrices were also tested by Adonis for true-differences in samples taken from different storm trajectories; no significance was detected. Snow filtering-process controls, DNA extraction controls, and no-template PCR controls (NTCs), amplified during qPCR a minimum of two cycles after at least one filter replicate from all samples, indicating that differences seen between samples are true biological differences. Filtering-process controls, furthermore, were ordinated against true samples and found to be distinct in clustering (data not shown).

**Figure 4:**
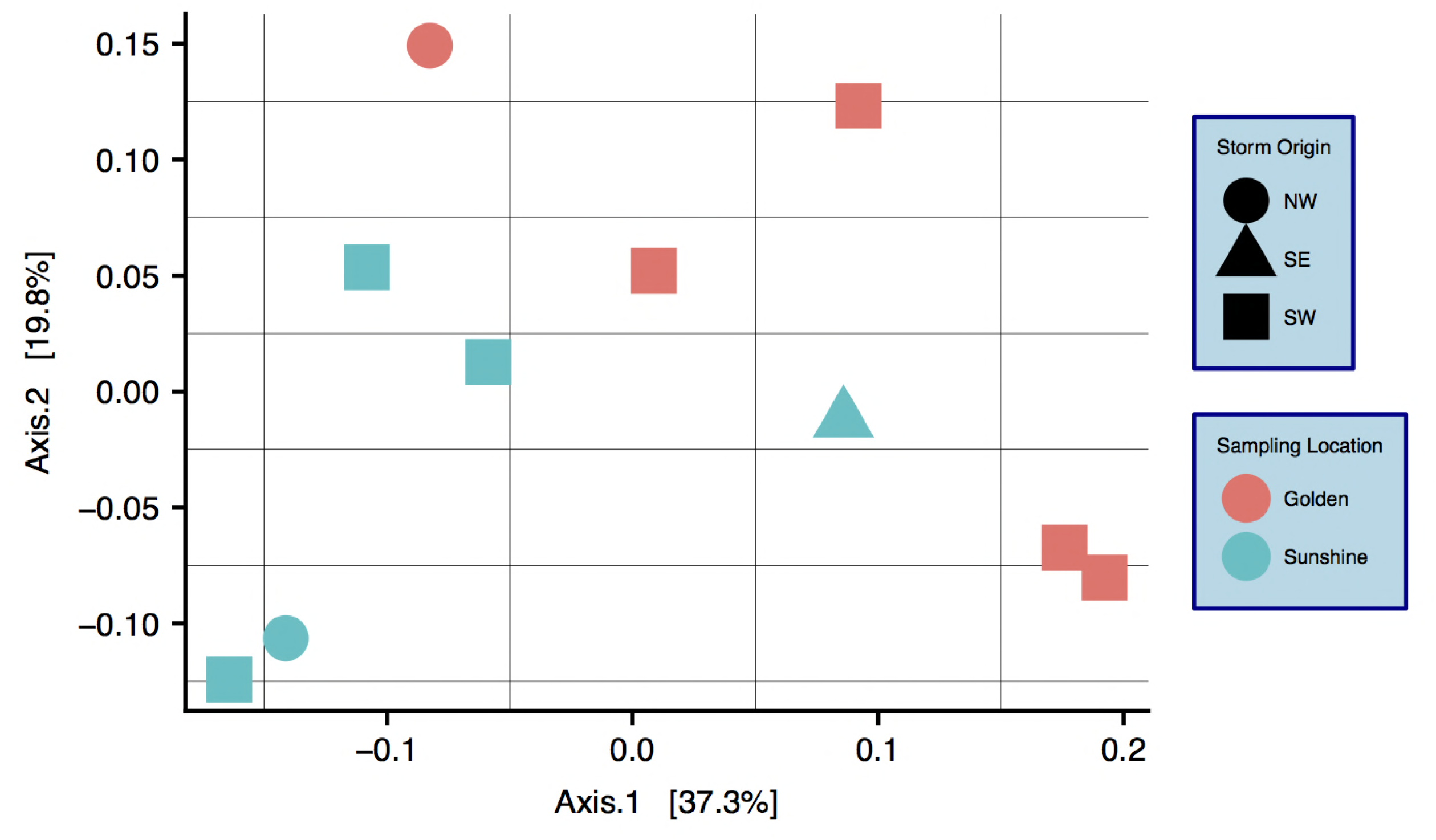
Principal Coordinate Analysis (PCoA) of a weighted Unifrac distance matrix. Colors are by Sampling Location and shapes are by Origin of Storm. Differences between samples with respect to Sampling Location are significant: R^2^ = 0.20, p = 0.033 (Adonis test).

Snowstorms from both sampling locations display a wide range of bacterial / archaeal (Figure S4A) and eukaryotic (Figure S4B) taxa. At the phylum level, *Bacteroidetes* are most common in snow sampled from Golden while *Proteobacteria* are most abundant in Sunshine samples, this in a linear distance difference of only 36.4 kilometers. At all taxonomic ranks, there are distinguishable differences in community composition between the two geographic locations. By use of SIMPER (vegan R package) (40), the taxa from each taxonomic rank most responsible for the Bray-Curtis distance between samples were determined. Plotted in descending order of contribution, we demonstrate how any taxonomic rank is sufficient to differentiate the two sampling locations (Figure 5). The top-ten most influential genera in differentiating sample locations were pulled from the SIMPER analysis, and are shown in Table 1 along with their % contribution to the total difference. The abundances of these genera are examined from both sampling locations as well as the control from the filtering process to assess contamination; no contamination is suspected (Figure S6). For each of the top ten most influential genera (Table 1), the OTU that had the highest abundance in each genus bin was searched with MegaBLAST (41) for highly similar sequences within the 16S rRNA sequence database from NCBI. For each sequence searched by MegaBLAST, the most highly similar sequence with information on source was summarized: genera found more frequently in Golden were associated with soil, horse dung, airborne dust, hard water, and clinical isolates (42–46); genera common to Sunshine are reported in swimming pool water, clinical material, cultivated land, and may be spore forming (47–51). Though Adonis showed no significant difference between samples with different storm-origins via the weighted Unifrac distance, there are obvious taxonomic differences between storm trajectories (Figure S5).

**Table 1:** The top-ten most influential genera in the biological differentiation of snowfall between geographic locations. SIMPER was used to determine the influence of each genus on the total Bray-Curtis difference between Sunshine and Golden samples. 279 unique genera were found in fresh snowfall.

| Genus | % Contribution to Distance | Location With Higher Abundance |
| --- | --- | --- |
| Chryseobacterium | 9.4 | Golden |
| Acinetobacter | 5.9 | Golden |
| Bacillus | 4.4 | Sunshine |
| Sphingomonas | 4.4 | Golden |
| Porphyrobacter | 3.2 | Sunshine |
| Flavisolibacter | 3.0 | Sunshine |
| Staphylococcus | 2.8 | Sunshine |
| Segetibacter | 2.7 | Sunshine |
| Pedobacter | 2.5 | Golden |
| Empedobacter | 2.3 | Golden |

**Figure 5:**
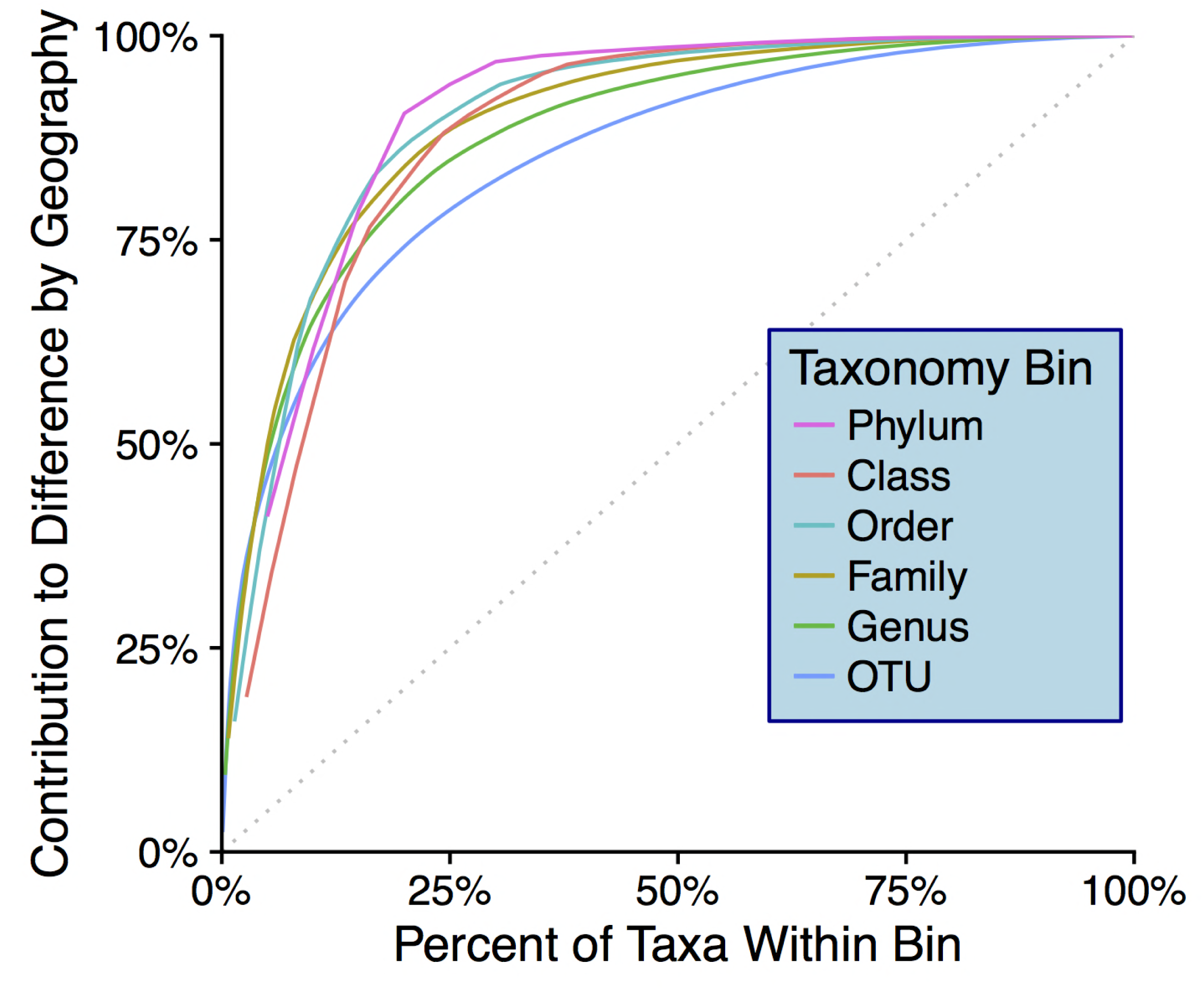
The y-axis shows cumulative contribution to the total Bray-Curtis distance between geographic locations. The x-axis indicates the top ‘X%’ most influential taxa within each bin that contribute to Bray-Curtis differences between geographic locations (determined via SIMPER analysis). The dotted gray line indicates what curves would look like if no particular taxa contributed to differences more than others.

Few eukaryotic sequences were recovered relative to the abundance of bacterial and archaeal sequences found in snowfall; prior to rarefaction, 6.2% of the total number of sequences recovered were of eukaryotic taxonomy. Of those eukaryotic sequences recovered and classified, the two fungal phyla *Ascomycota* and *Basidiomycota* were clearly most common. Similar to the results presented from bacterial and archaeal sequences, the two dominant eukaryotic phyla display differential relative abundance with respect to both sampling location and the origin of snowstorms (Figures S4B, S5B).

## Discussion

Snowfall is endemic to the seasonal cycle in both temperate and polar regions of the world. By contributing to aquifers, this form of precipitation is a critical component of both local and global hydrologic systems. Here we have presented a holistic approach to understanding the biology of meteoric snow, leveraging atmospheric phenomena as a means to determine significant affecters of biogeography. In our investigation, we have targeted the magnitude and community composition of meteoric snow as important first-steps in uncovering the importance of these fresh deposits to biogeography and the hydrologic cycle. Concerning magnitude, our data show that an average of 0.237 ng of bacterial / archaeal gDNA can be recovered per mL of fresh snowfall (melted and filtered), which is a value unreported in the literature. Our yields of DNA from snowfall are similar to, or greater than, what other groups have reported from snowpack (23, 24, 52), but we suggest that the annual persistence of meteoric snow coupled with massive land coverage yields a substantial amount of microbiota to surface ecosystems input.

These deposited microbiota also generate an influx of genetic material to receiving systems. The mean annual snowfall that occurred in Boulder, CO (NOAA, Boulder, CO) during 2010-2016 was 248.2 cm (1). This value, coupled with dimensional analysis, shows that given the mean bacterial / archaeal gDNA recovery that we report (0.237 ng mL^-1^ of snowmelt; ~4.7 × 10^4^ cells mL^-1^), a season-average conversion of snow volume to water volume (10:1, NOAA, Boulder, CO) (1), and a representative genome mass from *E. coli* (5 fg/cell), there are large biological depositions from snow computed for small areas: ~1 × 10^14^ bacterial / archaeal cells are deposited hectare^-1^ year^-1^. If one is to estimate the average mass of a bacterial / archaeal cell in our samples as 0.1 ng (a rough abundance-weighted average of the masses of bacterial / archaeal cells we identified taxonomically—including the large cyanobacteria), ~10 kg of bacteria / archaea are deposited hectare^-1^ year^-1^. Though this annual deposition of cells is calculated via multiple variables with internal variance (especially when estimating average cell mass), it is concomitantly noted that what we are able to recover through a 0.45-μm filter is likely an underestimation of what biological material exists. Microbiota captured on a filter, additionally, does not account for additional biomass that may have lysed during the thawing of our snow samples prior to filtering and/or the organisms, e.g., nano-archaeota and bacteria (as well as viruses) that pass through a 0.45-μm filter. Many eukaryotic sequences were also obtained from our samples. Though we do not have qPCR data on eukarya in fresh snow, this additional biomass (6.2% of all sequences)—known to be legitimate eukaryotic amplification by our primers (32)—is likely to contribute substantially to the total biomass contained in snowfall. Thus, 10 kg of biomass deposited hectare^-1^ year^-1^ is likely an underestimate. In addition, this estimate ignores the mass of all abiotic particulate matter (Figure 1) trafficked as part of the same snow event—a mass that is much more substantial than microbial elements alone and, similarly, is deposited perennially over large swaths of land.

Herein we have chosen to use the total estimated biomass of annual snowfall over more traditional measures like copy numbers of ssuRNA 16S because we believe that it is a more tractable unit when assessing large-scale mass transport and deposition. It is our assessment that the importance of our work is best demonstrated by the surprising bio-load of meteoric snow (and its associated genetic bank)—and that whether total estimated cell count or total biomass is the unit of discussion (both of which we report) is tangential to the overall finding that microbiota and their genomes are a substantial component of fresh snowfall.

Since we show significant differences between snowfall communities with respect to geography (Figure 4), we posit that snow depositions bear significant ecological consequences— especially given that we demonstrate the mass of any particular snow event to be non-trivial (Figure 3). At all taxonomic ranks, we show that just a fraction of taxa are responsible for differentiating geographic locations (Figure 5). To the point of ‘fingerprinting’ snowstorms to evaluate how different depositions may be affecting an area ecologically, it is important to identify specific taxonomic groups that are responsible for separations on distance matrices and ordinations (Table 1). We can infer from Table 1 that, even at the genus taxonomic rank, specific taxa contribute heavily to geographic differences in fresh snowfall. Furthermore, these genera are not contamination (Figure S6), indicating that real biological differences can be identified geographically in non-general terms. Of the influential genera, sequences that are more abundant in Golden (searched by MegaBLAST—Results section) are highly similar to some sequences that are associated with the human built environment / urban areas (i.e. horse dung and clinical isolates). Influential sequences from Sunshine, similarly, are highly similar to what is found in rural settings (genera isolated from cultivated land); one of the sequences more common in Sunshine is also highly similar to known spore-formers, suggesting that resilience to colder temperatures experienced in-transit to a higher elevation than Golden may have been beneficial to survival. Though we caution too much extrapolation from the presence of just a few genera, we think it is fair to suggest that demonstrated differences between Sunshine and Golden are the result of environmental / physical differences between the two geographic sites. Our control data, furthermore, strongly suggest that these differences are real, and not a product of contaminating genera found in either sample group (Figure S6). Not only do our data show that different geographical regions receive distinct biological input, but they also indicate that storm trajectories will be an essential metric to assess in future studies. Although we do not show demonstrable significance in the microbiological differences between storm trajectories, storm-track is still likely a contributing component of the variance between samples (Figure S5). Even with strong evidence that geography drives differences in microbial communities, we note that the two axes from our principal coordinate analysis represent less than 60% of the total variance seen between samples. Our investigation of storm-track influence on community composition is likely hindered by sample size; i.e. amongst the 10 samples profiled, only three of them were not of a SW origin. Overall, our hypothesis that the biotic components of fresh snowfall are distinguishable by geography is confirmed. To the point of meteorological information as predictive data in future studies, storm tracks should not be ignored amongst the list of affecters of microbial community composition. With ongoing collection of snow from more storms and winters, storm tracks could be forensically investigated by an analysis of their surface-deposited microbiota in snows, as well as the ices that result from snows in / on permanent snow fields and glaciers.

Bowers et al. showed in 2011 (7) that air mass microbiological features were likely derived from the land beneath. We extend this finding by showing that geography also appears to have community effects on snowfall deposition. We, however, further posit that biological ice-nucleation in the atmosphere (53, 54) may be a selection event influential enough to differentiate microbial communities even more than air masses do at any given geography. Thus, snowfall provides definitive access to surface receiving systems whereas air masses may not be as capable of moving large amounts of material between the atmosphere / lithosphere / hydrosphere. Irrespective of known high-temperature ice-nucleating (IN) bacteria like *Pseudomonas* genera, other bacterial communities may affect low-temperature cloud glaciation (55). If we assume that microbiota deposited by snowfall were either directly part of the nucleation process—or indirectly accumulated along the way—then snow depositions could be just a subset of the total aerosol microbial population. Aerosol microbiota and cloud thermodynamics are clearly linked given that different cloud chemistries determine what bacterial species act as nucleation points, and whether snow forms at all (56, 57). In light of our study, microbial communities should still be considered an important part of the total air mass that affects cloud formation potential and the precipitation nucleation process. Though it is known that high-temperature INs are ubiquitous in precipitation (58), our work demonstrates that the vast majority of microbiota in fresh snowfall are not of the high-temperature nucleating genera. More attention should be given to total bacterial communities in precipitation due to the magnitude of their abundance, potential for terrestrial impact, likely effects on cloud glaciation, and non-uniform patterns of deposition in snowstorm events (Figure 4). We know that bacteria/archaea/eucarya can be viable in aerosols (59), suggesting that the multitude of taxa that we find in snow may be contributing to atmospheric chemistry *in*-*situ* around Earth. Given all of the bio-physical / chemical influences on the atmosphere, bio-precipitation should be considered just as complex and variable—and storm-track should be considered and essential component of this variance. Furthermore, the importance of urban / rural areas receiving entirely different inputs in the same snowstorm cannot be understated, especially given the large mass of these annual deposits. This could have relevance to overall air quality and human health in such storm events.

Bio-aerosols are known components of human-associated dust particle traffic (12), and dust has complicated snowmelt and ecosystem function (60, 61). Our work shows that high annual flux of genetic material and biomass via snowstorms is non-homogenous. We can immediately suggest that human activity in urbanized areas could be a source of both the biological and geochemical differences witnessed in snow relative to rural mountain collection sites. Western regions of Colorado—the mountains—are less developed than the areas surrounding the greater Denver metropolitan area; i.e. the population density of Jefferson County (Golden samples) is roughly 70% greater than Boulder County (Sunshine samples) (62). Nonetheless, some similarities exist between the microbiology we found in fresh snowfall and taxa found by Amato et al. in the water phase of tropospheric clouds (20). Though several factors appear to influence the microbiological composition of snowstorms, a large contributor may be whether or not a certain bacterium is able to remain viable at low temperatures within a cloud, both low- and high-altitude. Community differences in snowfall between storm travel directions (Figure S5)— though insignificant due to our small number of samples from atypical storm trajectories—could also suggest that material collected along the path of a storm is later deposited as snow.

Broad questions on the impact of snowfall-deposited microbiology on both public health and ecosystem function can be generated by the work we describe herein. With respect to public health, in light of our work we believe that snowfall (and meteoric deposition of any kind –e.g., rain, snow, ice, fog and perhaps even general humidity) can be considered a large transporter of microbial biomass with a (to-date) not understood pathogenicity. In considering ecosystem function, the impact of some ~10 kg of bacterial / archaeal biomass being deposited per hectare in a year of snowfall clearly must include effects on receiving soil, rock endolith and/or near-surface water ecosystems. This has impacts on ecological soil formation, maintenance and health of agricultural soils and water quality in both fresh and marine ecosystems. As climate change continues to be a driver of large-scale environmental shifts, the intricacies of biological aerosol transport and bio-cloud glaciation / snow nucleation are unlikely to escape disturbance.

The flux in genetic diversity brought about by fresh snowfall is in much need of greater study. The amount of gDNA deposited by snow per hectare of land suggests that gene flow, via known mechanisms of horizontal gene transfer, is likely widely seeded by continual deposition from storm events. Ecosystem *in*-*situ* genetic diversity is thus likely to be in a state of higher flux than previously acknowledged, with receiving ecosystems likely more capable of responding to dynamic environmental conditions with updated ‘genetic banks’, literally always raining upon them. As we argue that the biomass within meteoric snow is ecologically impactful, we acknowledge that one may posit that what is deposited in snow would be just a fraction of the biomass already existing at the soil / hydrologic interface. We point out that meteoric snow would certainly be less consequential if it were uniformly distributed to all surface communities. However, microbiota within snow are highly heterogeneous (even within the same storm) and alternate genetic banks are deposited in different regions, some of which may be native to an area already—but many are likely to be non-local. Within oceanic systems, McDaniels et al. show that up to 47% of culturable bacteria are confirmed gene recipients by known mechanisms of horizontal gene transfer (63); if one year of snowfall biomass represents even 0.1% of the cells within the top 1 cm of receiving soil (assuming 1 × 10^9^ cells cm^-3^), those cells would be in sufficient quantity to impart new genetic material on the host community. If surface communities demonstrated even a fraction of the horizontal gene transfer ability of oceanic communities, a substantial portion of that entire 1 cm layer (and more) would be the beneficiary of new genetic material—accelerating genome innovation (64). Bacteria / archaea genomic innovation and resiliency has largely been driven by horizontal gene transfer (65) and we expect that environmental systems comprised of these organisms are likely to depend upon this mechanism for survival. Since lateral gene transfer increases genomic diversity at rates much higher than *in*-*situ* evolution alone (65), communities that receive higher fluxes of genetic material are likely to be more resilient than those that receive very little. We demonstrate in our investigation that geographic areas receive distinct biological input. This suggests that ‘genetic banks’ received by surfaces during snow storms send different locales on alternate genetic uptake and evolutionary pathways. Genetic transport by snowfall (and rain / ice / humidity) disallow microbial communities at any given geography to remain isolated. In an Earth system that is increasingly affected by climate change, the ability of local microbial communities to adapt will become paramount; atmospheric microbial transfer and snow / rain / icing / humidity events should be considered essential components in delivering novel genetic material to geographic areas. The ability for soil / water receiving microbial systems to remain resilient will have demonstrable effects on downstream ecological pathways such as macroscopic life and the mediation of soil / water chemical equilibrium—including the management of contaminants. The ‘everything is everywhere’ notion of microbial distribution across the globe can likely be better understood by the interaction of Earth’s compartments (lithosphere, hydrosphere and atmosphere) via transport. On-going work with metagenomics, transcriptomics and single-cell genomics should better inform the surface ecosystem-altering effects of the genetic potential of storm delivered events.

We suggest that bio-meteorology is likely to have impacts far beyond the Colorado Front Range alone; everything that we describe here is globally relevant to the lithosphere / hydrosphere / atmosphere interface in tropical, temperate and polar climates. Indeed, biology likely needs to be better incorporated into meteorology and meteorologic modeling. Disparate concepts ranging from ecological forest fire recovery to regional public health and pathogen transport are likely to be affected by the geochemical and biological depositions that we have described in snowfall (though these findings are likely also relevant to all meteoric waters including rain, fog, icing events and humidity). Further, biometeorology could likely be better informed and made capable of public-health related predictability with further microbiome understanding of widespread and large-scale meterologic events.

## Acknowledgements

We would like to thank all members of the GEM (Geo-Environmental-Microbiology) Lab at the Colorado School of Mines for helpful discussions. Specifically, thank you to Blake Stamps and Gary Vanzin for feedback on bioinformatic techniques and experimental design. We also would like to thank reviewers for helpful commentary that assisted in the improvement of our work.

## Figure Legends and Tables

Figure S1: Rarefaction curves generated by rarefying bacterial / archaeal sequences at different levels with random OTU sub-sampling. Each curve and color represents a different sample. A
blanket rarefaction value of 3,391 sequences was chosen for the bacterial and archaeal sequence dataset.

Figure S2: NOAA HYSPLIT storm trajectory model generated for the sample collection in Sunshine on 04/04/17. Streamlines are a medley of trajectory models distributed about the latitude and longitude of the sampling site. The backward trajectories encompass 24 hours of storm movement prior to arrival at the sampling location at an altitude of 500m AGL (above ground level). This storm was classified as having a NW origin. See Figure S3 for the Golden counterpart to this storm.

Figure S3: NOAA HYSPLIT storm trajectory model generated for the sample collection in Golden on 04/04/17. Streamlines are a medley of trajectory models distributed about the latitude and longitude of the sampling site. The backward trajectories encompass 24 hours of storm movement prior to arrival at the sampling location at an altitude of 500m AGL (above ground level). This storm was classified as having a NW origin. See Figure S2 for the Sunshine counterpart to this storm.

Figure S4: Relative abundances (more red = higher relative abundance) of bacterial / archaeal **(A)** and eukaryotic **(B)** sequences recovered from snow samples pooled by Sampling Location. Class-level assignments are listed first on the y-axis for each taxa, followed by a semi-colon and its associated phylum. Taxonomy was assigned via the SILVA database by DADA2. These data have been rarefied.

Figure S5: Relative abundances (more red = higher relative abundance) of bacterial / archaeal **(A)** and eukaryotic **(B)** sequences recovered from snow samples pooled by Storm Origin. Class-level assignments are listed first on the y-axis for each taxa, followed by a semi-colon and its associated phylum. Taxonomy was assigned via the SILVA database by DADA2. These data have been rarefied.

Figure S6: Demonstration that the most influential genera listed in Table 1 are not contamination. Numbers are relative abundances; more red means higher abundance. These data have been rarefied the same as all other analyzed samples.

